# A Single-copy Knock In Translating Ribosome ImmunoPrecipitation (SKI TRIP) tool kit for tissue specific profiling of actively translated mRNAs in *C. elegans*

**DOI:** 10.1101/2021.12.22.473890

**Authors:** Laura E. Wester, Anne Lanjuin, Emanuel H. W. Bruckisch, Maria C. Perez Matos, Caroline Heintz, Martin S. Denzel, William B. Mair

## Abstract

Translating Ribosome Affinity Purification (TRAP) methods have emerged as a powerful approach to profile actively translated transcripts in specific cell and tissue types. Epitope tagged ribosomal subunits are expressed in defined cell populations and used to pull down ribosomes and their associated mRNAs, providing a snapshot of cell type-specific translation occurring in that space and time. Current TRAP toolkits available to the *C. elegans* community have been built using multi-copy arrays, randomly integrated in the genome. Here we introduce a <u>S</u>ingle-copy <u>K</u>nock In <u>T</u>ranslating <u>R</u>ibosome Immuno<u>P</u>recipitation (SKI TRIP) tool kit, a collection of *C. elegans* strains engineered by CRISPR in which tissue specific expression of FLAG tagged ribosomal subunit protein RPL-22 is driven by cassettes present in single copy from defined sites in the genome. In depth characterization of the SKI TRIP strains and methodology shows that 3xFLAG tagged RPL-22 expressed from its endogenous locus or within defined cell types incorporates into actively translating ribosomes and can be used to efficiently and cleanly pull-down cell type specific transcripts without impacting overall mRNA translation or fitness of the animal. We propose SKI TRIP use for the study of processes that are acutely sensitive to changes in translation, such as aging.

## Introduction

Powerful methods have been developed to study translation and the translatome, the proportion of mRNAs that are actively undergoing translation. Polysome profiling monitors the efficiency of the translational machinery and can distinguish mRNAs translated by one ribosome (monosome) or several ribosomes at the same time (polysomes), and determine their relative rates of translation (Mašek et al., 2011). Ribosome footprinting maps ribosomes on the mRNA at codon resolution, allowing the analysis of translational speed as well as translational start and stall sites (Ingolia, 2016). Using these methods to analyze actively translated mRNAs within specific cell- or tissue-types however can be complicated by the presence of neighboring tissues and cells during the tissue isolation processes, especially in small model organisms such as *Drosophila* or *Caenorhabditis elegans* (*C. elegans*). Translating Ribosome Affinity Purification (TRAP) techniques elegantly overcome this problem and enable clean tissue- and cell-type specific investigation of the translatome. Here, ribosomal proteins are tagged and used for co-immunoprecipitation (co-IP) of fully assembled ribosomes and their associated mRNAs (King and Gerber, 2016). Expression of tagged ribosomes selectively within genetically defined cell types can be used to provide a snapshot of their cell type-specific translatome and identify the changes occurring upon treatment or over time.

Integral to the TRAP technique is the addition of an epitope tag to a ribosomal subunit protein to use for co-immunoprecipitation (co IP). To date, all existing TRAP models have tagged a protein of the 60S large ribosomal subunit and expressed it tissue- or cell-type-specifically. The first animal TRAP models were successfully generated in mice by tagging RPL10a (Heiman et al., 2008) or RPL22 (Sanz et al., 2009), which sit at the surface of the assembled 80S ribosome. Similar approaches have been adapted to *Drosophila* (Desplan et al., 2017; Thomas et al., 2012) and *C. elegans* (Gracida and Calarco, 2017; McLachlan and Flavell, 2019; Rhoades et al., 2019). Studies conducted with these genetic models have used TRAP to profile translated mRNAs in genetically defined cell types or under distinct physiological conditions. In mice, TRAP has facilitated the molecular characterization of intermingled neuron subtypes in the brain (Chandra et al., 2015; Heiman et al., 2008). In *C. elegans*, TRAP has been used to profile tissue-specificity in alternative splicing (Koterniak et al., 2020), and to identify genes enriched in serotonergic neurosecretory motor neurons that regulate food sensing and stress-induced feeding (Gracida et al., 2017; Rhoades et al., 2019).

Although TRAP has proven to be an extremely powerful tool for characterizing genetically defined cell types and the differences between them, applying TRAP approaches for the study of processes such as aging, metabolism and stress response pathways that are acutely affected by changes in translation itself poses a unique set of challenges. Most important is that the TRAP tools themselves must not cause changes to translation. For example, the presence of the epitope tag on the ribosomal subunit must not alter ribosomal subunit function in any way leading to impaired translation, as changes of translation *per se* can for example alter the lifespan of *C. elegans* and higher organisms (Hansen et al., 2007; Pan et al., 2007; Syntichaki et al., 2007). Further, for use of the TRAP approach for the study of tissue-specific changes to translation itself, it is important that the epitope tag on the ribosomal subunit not result in changes to translation rate, ribosomal distribution, or lead to biases in transcript usage.

All current TRAP models developed in *C. elegans* have been made using integrated extrachromosomal arrays, which are commonly used to generate tissue-specific gene expression (Gracida and Calarco, 2017; McLachlan and Flavell, 2019; Rhoades et al., 2019). Extrachromosomal arrays are high copy number, repetitive transgenes that often result in extremely high levels of expression. The consequences of altered levels of a tagged ribosomal protein to translation-sensitive processes such as aging is not clear. In addition, integration of these extrachromosomal arrays into the genome via random mutagenesis insertion can disrupt coding or regulatory sequences. Further, where linkage prevents crossing randomly inserted TRAP cassettes into other genetic mutants, it is impossible to recreate identical copy number cassettes inserted at alternate loci. To adapt the TRAP approach for studies where changes in translation need to be minimal and well controlled, we set out to generate and fully characterize a *C. elegans* TRAP tool kit by CRISPR in which tissue specific transgenes are present in single copy, and the impact on translation and to the general fitness of the animal is known.

Here we present a <u>S</u>ingle-copy <u>K</u>nock In <u>T</u>ranslating <u>R</u>ibosome Immuno<u>P</u>recipitation (SKI TRIP) tool kit, a collection of strains in which an N-terminal 3xFLAG tagged version of the conserved 60S ribosomal subunit protein RPL-22 is expressed across a panel of tissue types from defined intergenic sites in the genome in single copy. We show by expressing FLAG-tagged RPL-22 from its endogenous locus that the presence of the FLAG tag has no impact on overall translation and does not result in any bias in actively translated transcripts. FLAG-tagged RPL-22 expressed from its endogenous locus or within defined cell types incorporates into actively translating ribosomes without impacting overall mRNA translation or fitness of the animal, and can be used to efficiently and cleanly pull down cell-type specific transcripts. We introduce the SKI TRIP tool kit and provide a comprehensive analysis of the cell type specific translational state captured by each of the strains.

## Results

### Design and generation FLAG tagged RPL-22 *C. elegans* strains

With the aim of generating a *C. elegans* SKI TRIP tool kit that expresses a tagged ribosomal subunit from defined loci using well-characterized expression cassettes present in single copy, we first sought to determine the consequences of tagging the subunit on overall ribosomal function. In analogy to published TRAP models, we opted to tag RPL-22, a protein of the 60S large subunit of the *C. elegans* ribosome (Fig. 1A) (Rhoades et al., 2019; Sanz et al., 2009). We generated knock-in alleles of the endogenous *rpl-22* gene, *rpl-22(wbm58) and rpl-22(wbm53)*, that introduce 3xFLAG sequences either onto the 5’ or 3’ end, respectively (Sup. Fig. 1A and B). Both alleles are viable and fertile, suggesting either the N- or C-terminus of RPL-22 can support a FLAG tag without grossly impairing its function (data not shown). GFP knock-in to the 3’ end of *rpl-22* confirms that this ribosomal subunit is robustly and ubiquitously expressed (Fig 1B). In addition, expression of *rpl-22* is stable with age, as characterized by RNAseq analysis (Heintz et al., 2017) (Sup. Fig. 1C). Interestingly, homozygous *rpl-22(wbm85)* GFP knock-in animals are slow growing and sterile, suggesting incorporation of the larger GFP tag as opposed to the smaller FLAG tag on the C-terminus is likely detrimental to RPL-22 function (data not shown).

**Figure 1:**
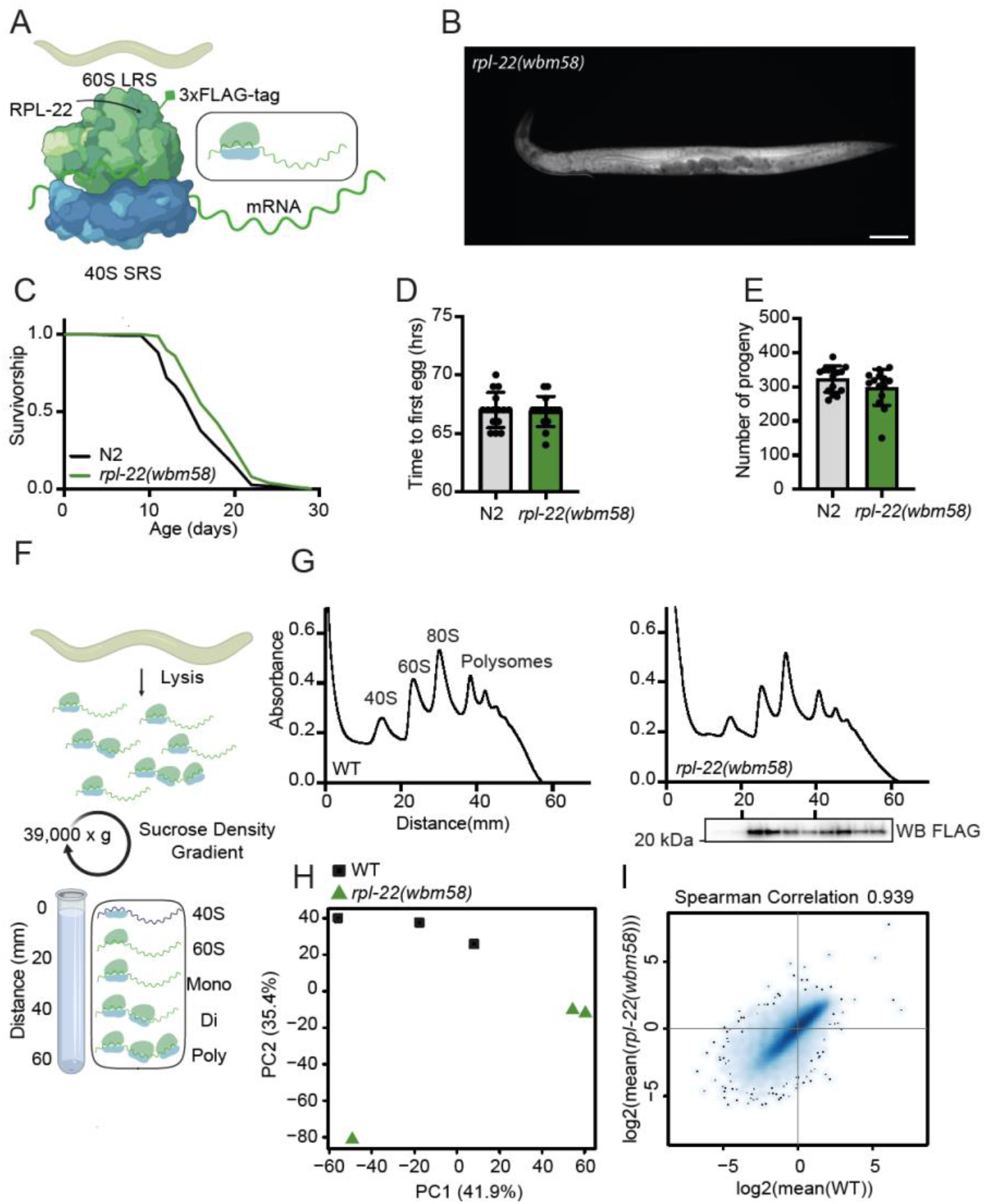
N-terminal FLAG-tag on RPL-22 does not impact *C. elegans* physiology or overall mRNA translation. (**A**) Model of a fully assembled ribosome including associated mRNA between the small ribosomal subunit (40 S) and the large ribosomal subunit (60 S) showing FLAG-tagged RPL-22 at the ribosomal surface. (**B**) Fluorescence microscopy image of a heterozygous *rpl-22(wbm85)* (RPL-22::GFP knock-in allele) animal at Day 1 of adulthood. Scale bar = 100 µm. (**C**) Survival of *C. elegans* carrying an endogenous, C-terminal FLAG-tag at RPL-22 (endogenous SKI TRIP strain *rpl-22(wbm58)*), compared to WT (N2) (representative data from n=3 independent experiments). (**D**) Generation time of WT and *rpl-22(wbm58)* animals (error bars represent means ±SD; representative data from n=3 independent experiments with 15 animals per genotype). (**E**) Brood size of WT and *rpl-22(wbm58)* as total number of viable progeny per individual parental animal (error bars represent means ±SD; representative data from n=3 independent experiments with ≥ 12 animals per genotype). (**F**) Schematic depiction of the polysome profiling approach. After lysis, ribosomal subunits, monosomes (80 S) and polysomes are separated by centrifugation on a sucrose gradient. (**G**) Polysome profiling graphs from WT and *rpl-22(wbm58)* samples obtained at Day 1 of adulthood. Western blot analysis of extracted protein was performed from polysome profiling fractions of endogenous SKI TRIP worms as indicated in the figure. Anti-FLAG antibodies were used. (**H** and **I**) After polysome profiling, RNA was extracted from actively translated polysome fractions (see **G**) and pooled per strain. RNA was sequenced and values were normalized to total RNA each from WT and *rpl-22(wbm58)* samples (n=3 independent experiments were performed). (**H**) PCA comparing WT and *rpl-22(wbm58)* samples, using ratios of RNA expression values (FLAG IP RNA/(total RNA + 1)) from n=3 independent experiments. PC = Principal Component. (**I**) Scatter plot showing mean log2(ratios) of RNAs commonly detected in WT and *rpl-22(wbm58)* samples from n=3 independent experiments. See Sup. Figure 1 for detailed information on SKI TRIP strains. See Sup. Table 1 for lifespan statistics. See Sup. Table 3 for differential gene expression tables corresponding to the RNA sequencing experiments.

### Characterization of *C. elegans* with an endogenous ^FLAG^RPL-22 tag

To determine whether the presence of the N-terminal 3xFLAG tag on RPL-22 interferes with ribosome assembly or translation, or on its incorporation into actively translating ribosomes, we characterized our endogenous ^FLAG^RPL-22 knock-in allele, *rpl-22(wbm58)*, in which the endogenous *rpl-22* gene has been edited and all RPL-22 molecules are tagged (endogenous SKI TRIP animals). *rpl-22(wbm58)* animals exhibit no pronounced changes to lifespan, generation time or brood size (Fig. 1C-E) suggesting the presence of the N-terminal FLAG tag does not impair overall fitness of the animal. To look at the effect of N-terminally 3xFLAG tagging RPL-22 on translation specifically, we compared polysome profiling of wildtype (WT) and *rpl-22(wbm58)* lysates after separation on a sucrose gradient (Fig. 1F). *rpl-22(wbm58)* polysome profiles resemble those of WT, indicating functional translation machinery in ^FLAG^RPL-22 animals (Fig. 1G). Further, Western blot analysis of protein extracted from the sucrose gradient fractions shows that the FLAG epitope can be detected in the fractions of the 60S large ribosomal subunit, monosomes and in polysomal fractions, indicating ^FLAG^RPL-22 is incorporated into actively translating ribosomes (Fig. 1G). Taken together, these results suggest that tagging RPL-22 does not disturb normal ribosomal function or overall physiology, and that ^FLAG^RPL-22 can be incorporated in translating ribosomes in the monosome and polysome states.

To determine whether the presence of the FLAG tag on ^FLAG^RPL-22 biases the mRNAs that ribosomes associate with, we captured all actively translated mRNAs in WT and in *rpl-22(wbm58)* animals by polysome profiling and analyzed them by RNA sequencing (RNAseq) (n=3 per condition). Translated RNAs captured from each sample were normalized to total RNA, and then then tested bioinformatically for sample similarity (see methods). While WT and *rpl-22(wbm58)* mutant samples can be separated in a Principal Component (PC) analysis (Fig. 1H), their expression shows a high correlation (R_spearman = 0.939), indicating that their translatomes are highly similar (Fig. 1I). Taken together, these data suggest that the tagged ^FLAG^RPL-22 worms cannot be distinguished from WT animals by their polysome profiles or by their actively translated mRNAs.

### Immunoprecipitation of ribosomes from endogenous ^FLAG^RPL-22 animals

The principle of the TRAP approach is to use tagged ^FLAG^RPL-22 to pull down fully assembled ribosomes along with their associated mRNA (Fig. 2A). Western blot analysis after IP confirms that FLAG-tagged RPL-22 can be pulled down from *rpl-22(wbm58)* samples but not those from WT (Fig. 2B). Blotting for RPS-6, a protein of the small ribosomal subunit, shows that it too can be detected in samples from *rpl-22(wbm58)*, and therefore the IP pulls down assembled ribosomes. Lastly, we extracted RNA captured by *rpl-22(wbm58)* IP. By tape station analysis both the 18S and 28S rRNA species are detected and remain undegraded after ^FLAG^RPL-22 pulldown. The high RNA integrity number equivalent (RIN^e^) confirms that the pulled down RNA is of high quality (Fig. 2C).

**Figure 2:**
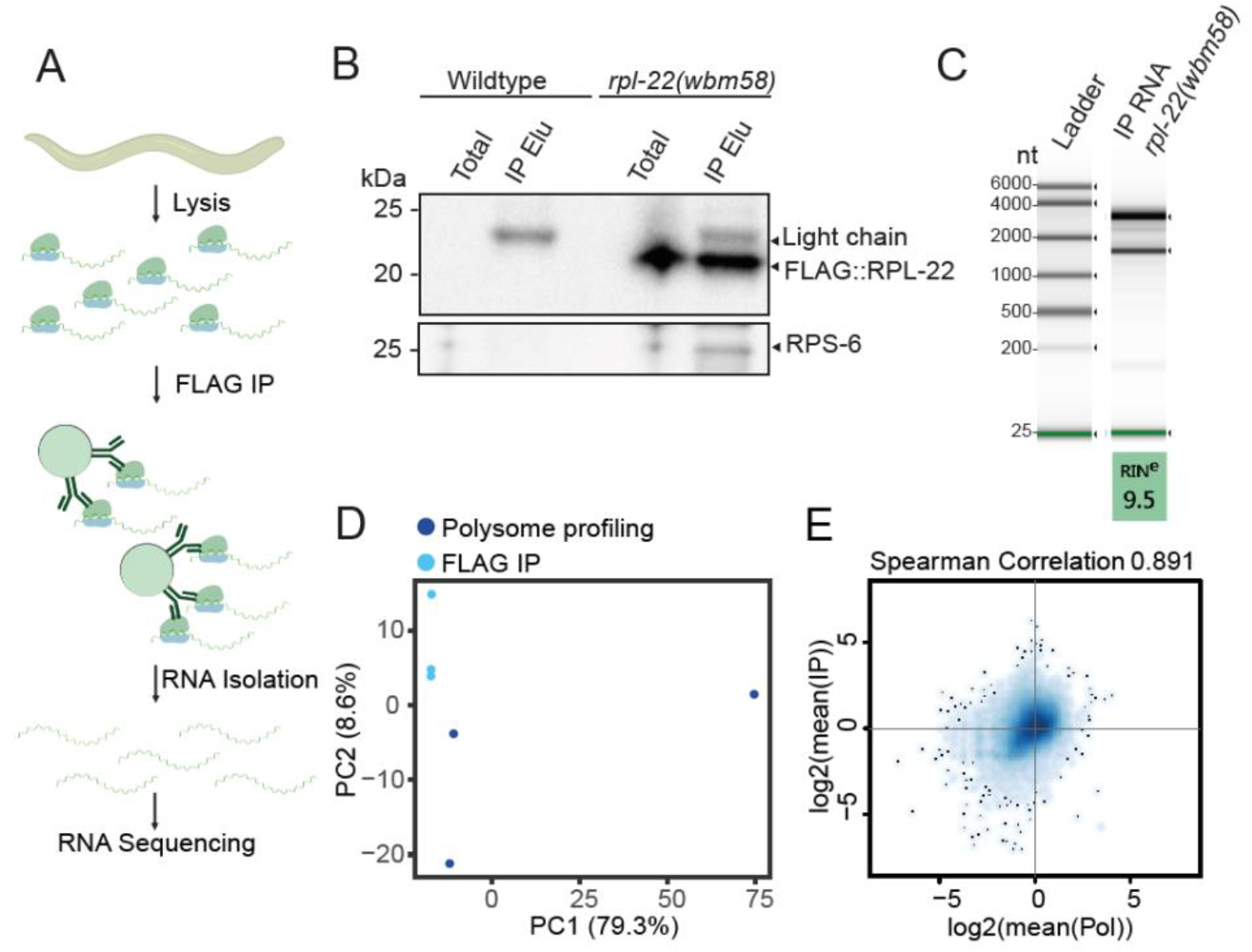
FLAG co-IP pulls down assembled ribosomes and associated RNA from endogenous SKI TRIP knock-in strain. (**A**) Schematic depiction of the FLAG co-IP approach using FLAG antibodies coupled to magnetic beads. (**B**) Western blot of day 1 WT and *rpl-22(wbm58)* samples after FLAG co-IP. Western blot using anti-FLAG and anti RPS-6 antibodies. co-IP total and elution (IP elu) fractions are shown. (**C**) On-chip electrophoresis of eluted RNA after FLAG co-IP from *rpl-22(wbm58)* samples, including RINe value. (**D** and **E**) RNA sequencing was performed from *rpl-22(wbm58)* samples: (1) with RNA captured from pooled 60S, 80S, monosomal and polysomal profiling fractions and (2) with RNA eluted after FLAG co-IP. Expression values were normalized to respective total RNA samples from each approach (n=3 independent experiments were performed). (**D**) PCA using mean ratios of RNA expression values (Eluted RNA/(total RNA + 1)) from n=3 independent experiments, comparing RNA from polysome profiling to RNA from FLAG co-IPs of *rpl-22(wbm58)* samples. PC = Principal Component. (**E**) Scatter plot showing mean log2(ratios) from n=3 independent experiments of commonly detected RNA in both polysome profiling and SKI TRIP IP samples from *rpl-22(wbm58)*. See Sup. Figure 1 for detailed information on SKI TRIP strains. See Sup. Table 3 for differential gene expression tables corresponding to the RNA sequencing experiments.

By all measures, the co-IP approach using the endogenous N-terminal FLAG tag knock-in allele demonstrates that the endogenously tagged *rpl-22(wbm58)* strain can be used to successfully profile actively translated mRNAs across all tissues. To determine whether RNA captured by ^FLAG^RPL-22 IP differs from RNA obtained by traditional polysome capture, we tested both methods back-to-back. We performed a polysome profiling experiment and compared the actively translated RNAs identified by RNAseq to those captured in an experiment using *rpl-22(wbm58)* IP. Despite the dramatically different experimental methods, by PCA analysis and by Spearman Correlation we found only small differences in the actively translated mRNAs identified by the two approaches (Fig. 2D, 2E, Sup. Fig. 2A). Whether these differences are due to variances in sample preparation methods or reflecting true biological differences between ribosomes containing RPL-22 and those as a whole is not yet clear. Taken together, these data show that the TRAP methodology using N-terminal ^FLAG^RPL-22 subunit expressed in all tissues can be used to generate a profile of actively translated mRNAs in the animal.

### SKI TRIP tool kit to analyze tissue specific translatomes in *C. elegans*

Reassured that the presence of the FLAG tag on RPL-22 does not negatively impact ribosomal function and that it can be used to efficiently pull down ribosome-associated RNA, we used the *C. elegans* <u>S</u>ingle-copy <u>K</u>nock-In <u>LO</u>ci for <u>D</u>efined <u>G</u>ene <u>E</u>xpression (SKI LODGE) system (Silva-Garcia et al., 2019) to engineer a collection of tissue-specific ^FLAG^RPL-22 strains by CRISPR: a ‘<u>S</u>ingle-copy <u>K</u>nock In <u>T</u>ranslating <u>R</u>ibosome Immuno<u>P</u>recipitation (SKI TRIP)’ tool kit. FLAG tagged *rpl-22* cDNA was inserted in single copy into well characterized tissue-specific SKI LODGE cassettes that are located in intergenic regions known to minimize any disruption to endogenous gene expression. As a starting point we generated strains for somatic, muscle, intestinal, and neuronal expression of ^FLAG^RPL-22, using cassettes driven by cell type-specific promoter sequences from the *eft-3, myo-3, nep-17, and rab-3* genes, respectively. The genomic constructs consist of the tissue-specific promotor followed by sequences for an N-terminal 3xFLAG tag, full-length *rpl-22* cDNA, a spliced leader (SL) RNA coupled with a wrmScarlet fluorophore and a *rab-3* or *unc-54* 3’UTR, depending on the cassette (full strain details can be found in Sup. Fig. 1A and B). Tissue specific expression of each strain is easily identifiable by fluorescence of wrmScarlet (Fig. 3A and Sup. Fig. 3A).

**Figure 3:**
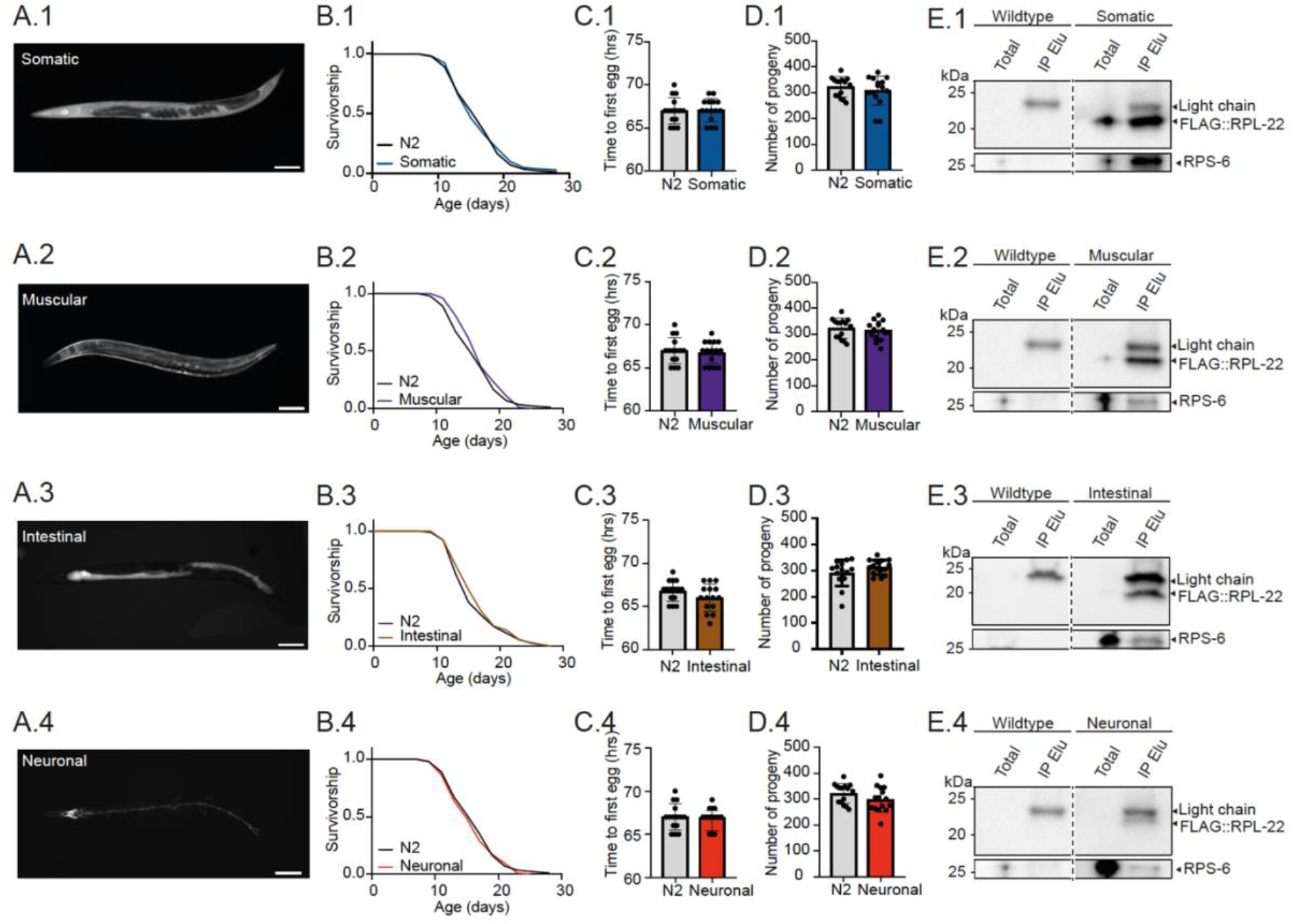
Tissue-specific expression of ^FLAG^RPL-22 in *C. elegans* SKI TRIP strains does not impact overall physiology and is sufficient for FLAG co-IPs. (**A.1-E.1**) Analysis of somatic SKI TRIP animals (*wbmIs119*). FLAG-tagged RPL-22 is expressed under the soma-specific *eft-3* promotor. The expression cassette is carrying a wrmScarlet fluorophore sequence after the FLAG::RPL-22; an SL2 sequence induces trans-splicing of the fluorophore from FLAG::RPL-22. (**A.1**) Fluorescence microscopy image of a day 1 somatic SKI TRIP animal. Scale bar = 100 µm. (**B.1**) Survival of somatic SKI TRIP animals compared to WT. (**C.1**) Generation time of WT and somatic SKI TRIP animals. (**D.1**) Brood size of WT and somatic SKI TRIP strain as total number of viable progeny per individual parental animal. (**E.1**) Western blot of day 1 WT and somatic SKI TRIP samples after FLAG co-IP. Western blot using anti-FLAG and anti RPS-6 antibodies. co-IP total and elution (IP elu) fractions are shown. (**A.2-E.2**) Analysis of muscular SKI TRIP animals (*wbmIs118*) as described above. FLAG-tagged RPL-22 is expressed under the muscle-specific *myo-3* promotor. (**A.3-E.3**) Analysis of intestinal SKI TRIP animals (*wbmIs131*) as described above. FLAG-tagged RPL-22 is expressed under the intestine-specific *nep-17* promotor. (**A.4-E.4**) Analysis of neuronal SKI TRIP animals (*wbmIs99*) as described above. FLAG-tagged RPL-22 is expressed under the neuron-specific *rab-3* promotor. (**A-E**) “Somatic” = soma-specific SKI TRIP *wbmIs119* [WBM1364] animals. “Muscular” = muscle-specific SKI TRIP *wbmIs118* [WBM1339] animals. “Intestinal” = intestine-specific SKI TRIP *wbmIs131* [WBM1470] animals. “Neuronal” = neuron-specific SKI TRIP *wbmIs99* [WBM1340] animals. Lifespan assays: Representative data from n≥2 independent experiments. Generation time and brood size assays: Representative data from n≥2 independent experiments with ≥ 13 animals per genotype; error bars represent means ±SD. See Sup. Figure 1 for detailed information on SKI TRIP strains. See Sup. Table 1 for lifespan statistics.

Based on the analysis of the endogenous ^FLAG^RPL-22 worms, we speculated that the tissue-specific expression of ^FLAG^RPL-22 should not interfere with general translation. To confirm this, we performed a broad analysis of worm health and translation on the SKI TRIP strains. Again, no differences in lifespan, generation time or brood size were observed in any of the strains compared to WT animals (Fig 3B, 3C and 3D and Sup. Fig. 3B, 3C and 3D). Polysome profiles performed on the somatic (*wbmIs119*), muscular (*wbmIs118*) and neuronal (*wbmIs99*) SKI TRIP strains are also indistinguishable from WT (Sup. Fig. 3F), indicating no effect of the tissue specific expression of ^FLAG^RPL-22 on overall translation. To test whether FLAG-IP can immunoprecipitate assembled ribosomes in sufficient amounts and quality from the tissue specific ^FLAG^RPL-22 single copy cassettes, we performed Western blot analyses after IP. We detected ^FLAG^RPL-22 and untagged RPS-6 in the elution fraction of samples from each of the SKI TRIP collection strains, but not from WT (Fig. 3E and Sup. Fig. 3E). Despite the more modest levels of expression of ^FLAG^RPL-22 in some of the newly generated tissue-specific strains, RNA obtained by co-IP from all strains in the SKI TRIP collection can be extracted in amounts and quality suitable for RNAseq.

Finally, to obtain a global picture of the mRNAs captured by IP from each of the respective SKI TRIP strains and to determine the extent to which they are enriched for cell type appropriate transcripts, SKI TRIP experiments were performed in three replicates each on samples of the somatic (*wbmIs119*), muscle (*wbmIs118*), and intestinal (*wbmIs131*) SKI TRIP strains, as well as from two neuronal strains (*wbmIs99, wbmIs133*) containing neuronal SKI TRIP cassettes inserted on different chromosomes. Extracted total and IPed RNA from each strain was sequenced. PCA analysis of all RNAseq experiments combined shows clustering of total RNA samples, reflecting overall high similarity of whole-body transcriptomes in the different strains, with defined separation of IPed samples, each clustering by tissue type (Fig 4A). In this analysis, IPed samples from the two neuronal lines cluster closely but separately, possibly due to their different sample preparation batches or to a change in RNA sequencing parameters (see methods). For further analyses, IPed RNA from each sample was normalized to its total RNA. Normalization was calculated either as ratios for the following heat maps, or as differential gene expression for the GO term analyses. To determine the nature of RNAs pulled down by each SKI TRIP, we manually compiled a list of cell type-specific genes from the literature and looked for their enrichment by IP. Heat map shows dramatic enrichment of cell type appropriate genes in IP samples from the tissue specific SKI TRIP strains (Fig 4B and Sup. Fig. 4A). GO analysis extends this analysis to a genome-wide level, showing significant upregulation of GO terms related to muscle, intestinal or nervous system function represented in the muscle, intestinal and neuronal SKI TRIP IPs, respectively, as well as a dramatic de-enrichment of gene classes implicated in embryonic development and reproduction characteristic of germline expression in IPs from the somatic SKI TRIP strain (Fig. 4C-F and Sup. Fig. 4B).

**Figure 4:**
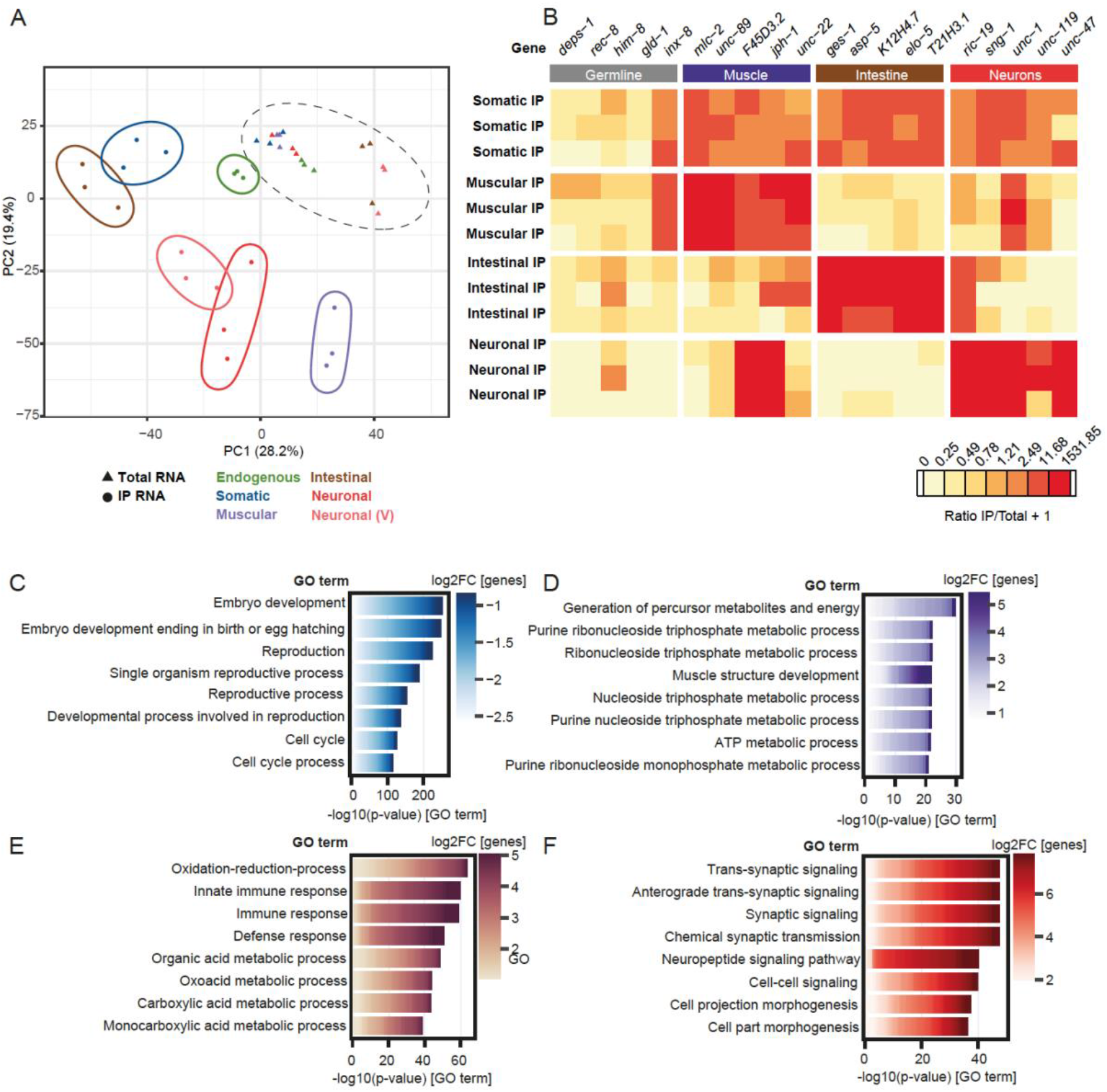
FLAG co-IPs from SKI TRIP strains cleanly pull down cell type specific mRNAs. FLAG IPs were performed from tissue-specific SKI TRIP worm strains in n=3 independent replicates. Total and elution RNA were sequenced. (**A**) PCA using batch-corrected RNA expression values, comparing total and elution RNA from FLAG co-IPs of endogenous, soma-, muscle-, intestine- and neuron-specific SKI TRIP samples. The 20 % most variable genes were used for the analysis. Data corresponding to three independent replicates from each strain are shown. While the original neuronal SKI TRIP strain carries the expression cassette on chromosome IV, a second strain *wbmIs133* (Neuronal (V)) was generated, carrying the expression cassette on chromosome V. Triangles represent co-IP total RNA, dots represent co-IP elution RNA. Solid ellipses comprise respective tissue-specific IP RNA samples, dashed ellipse comprises all total RNA samples. PC = Principal Component. (**B**) Heat map is showing expression values of FLAG co-IP elution RNA samples that were normalized to respective total RNA samples from each tissue-specific SKI TRIP strain. Data shown for selected genes that are known to be expressed in specific tissues as indicated in the figure. According to Wormbase (WS282), the genes *F45D3.2, jph-1* and *unc-1* are reported to also be expressed in muscle and neurons. The gene *inx-8* is reported to be expressed in germline and muscle. (**C**) DAVID GO term analysis of mRNAs significantly changed in translation (IP elution RNA values normalized to respective total RNA values of the same strain, using data of n=3 independent replicates). Analyzed were significantly down-regulated transcripts from the somatic SKI TRIP strain. Shown are the top 8 significant GO terms. (**D-F**) DAVID GO term analysis of mRNAs significantly changed in translation as in (**C**). Analyzed were significantly up-regulated transcripts from the muscular (**D**), intestinal (**E**) and neuronal (**F**) SKI TRIP strains. Shown are the top 8 significant GO terms. “Somatic” = soma-specific SKI TRIP *wbmIs119* [WBM1364] animals. “Muscular” = muscle-specific SKI TRIP *wbmIs118* [WBM1339] animals. “Intestinal” = intestine-specific SKI TRIP *wbmIs131* [WBM1470] animals. “Neuronal” = neuron-specific SKI TRIP *wbmIs99* [MSD473] animals, carrying the expression cassette on chromosome IV. “Neuronal (V)” = neuron-specific SKI TRIP *wbmIs133* [WBM1471] animals, carrying the expression cassette on chromosome V. See Sup. Figure 1 for detailed information on SKI TRIP strains. See Sup. Table 4 for differential gene expression tables corresponding to the RNA sequencing experiments. See Sup. Table 5 and 6 for full DAVID GO term analysis results.

Together these results show that single copy knock-in of FLAG tagged RPL-22 in the tissue specific SKI TRIP cassettes can be used to efficiently and cleanly profile the state of active translation within specific cell types. Neither expression of ^FLAG^RPL-22 nor the presence of the SKI TRIP cassettes themselves result in any observable changes to fitness or translation, suggesting this tool kit will be especially powerful for the study of mutants or conditions that are sensitive to changes in overall mRNA translation.

## Discussion

Here we introduce a new SKI TRIP tool kit to study tissue specific translation in *C. elegans*, engineered using the CRISPR/Cas9 based SKI LODGE system. We generated a panel of new *C. elegans* strains that expresses the 3x-FLAG tagged ribosomal subunit RPL-22 either endogenously or tissue specifically. We characterized in depth the effects of FLAG tagging RPL-22, ensuring the tag does not interfere with mRNA translation. We show co-immunoprecipitation of assembled ribosomes and associated mRNAs from ^FLAG^RPL-22. Finally, by RNAseq, we demonstrate dramatic enrichment of tissue-type appropriate mRNAs captured by SKI TRIP experiments.

Similar models have been developed for use in *C. elegans*. Gracida and Calarco (2017) showed that tagging the subunit RPL-1 of the *C. elegans* ribosome with enhanced green fluorescent protein (eGFP) can be used to isolate actively translated mRNA from body-wall muscle, intestine and the nervous system, including dopaminergic and serotonergic neuronal subtypes (Gracida and Calarco, 2017). Similarly, McLachlan and Flavell (2019), added an HA-tag to RPL-22 and expressed it specifically in serotonergic neurons (McLachlan and Flavell, 2019; Rhoades et al., 2019). Our tool kit compliments these existing models and provides some important advantages. We included an endogenously tagged RPL-22 strain *rpl-22(wbm58)* to our SKI TRIP tool box, that allows for tight control of the effect of FLAG-tagging RPL-22 on global mRNA translation. Further, in contrast to integrated extrachromosomal array-based expression where multi-copy transgenes are randomly inserted into unknown sites in the genome, SKI TRIP cassettes are present in single copy at known intergenic loci, with low overall expression levels (Silva-Garcia et al., 2019; Zhang et al., 2019). In addition, our model uses a 3xFLAG epitope instead of a substantially larger eGFP to minimize any effect of the tag on ribosome functionality. Indeed, we have shown that expression of ^FLAG^RPL-22 either tissue specifically from single copy cassette or from its endogenous locus has no observable effect on translation or on fitness of the whole animal, which was not the case when we tagged endogenous RPL-22 with GFP. Though RPL-22 itself is not fluorescently tagged in this tool kit, the design of the tissue-specific SKI TRIP cassettes allows the transgene to be followed via fluorescence of wrmScarlet. Lastly, our protocol decreases the amount of tissue needed for a pan-neuronal mRNA from over 150,000 worms (McLachlan and Flavell, 2019) to 30,000 worms and less, leading to a drastic reduction in consumables required for the IP reaction.

In the future, applying ribosome footprinting to tissue-specific ribosomes isolated from SKI TRIP worms will enable a detailed analysis of translational control with subcodon resolution (Ingolia et al., 2009). SKI TRIP experiments combined with polysome profiling on a sucrose gradient will make it possible to analyze the distribution of monosomes and polysomes on cell type-specific mRNAs (Ding and Großhans, 2009). It has recently been shown that ribosomes enriched with specific proteins (such as RPL-10A) have a higher likelihood of translating a subset of mRNAs that contain internal ribosome entry sites (IRES) (Shi et al., 2017), and that mRNAs with a certain Kozak sequence preferentially associate with ribosomes containing protein 26 of the small ribosomal subunit (RPS-26) (Ferretti et al., 2017). SKI TRIP isolation of ribosomes can be used to study such differences in ribosomal heterogeneity across different tissue types. All in all, several directions highlight the potential of the tissue-specific research on translation with the SKI TRIP method.

Here, we generate, characterize and distribute the SKI TRIP tool kit, an optimized and deeply characterized model to study tissue-specific translation for the *C. elegans* research community. Our in-depth demonstration that the N-terminal FLAG-tag at RPL-22 does not cause any observable effects on translation or fitness of the animals makes the SKI TRIP tool kit especially well-suited to study processes that are acutely sensitive to changes in translation, such as aging.

## Materials and methods

### Supplementary tables

All supplementary tables can be provided upon request.

**Supplementary Table 1:** Statistics on lifespan assays.

**Supplementary Table 2:** *C. elegans* strains used in this study.

**Supplementary Table 3:** RNAseq analysis results (DESeq2) corresponding to data shown in Fig. 1, Fig. 2 and Sup. Fig. 2.

**Supplementary Table 4:** RNAseq analysis results (DESeq2) corresponding to data shown in Fig. 4 and Sup. Fig. 4.

**Supplementary Table 5:** Full DAVID GO term analysis results based on down-regulated differential gene expression values of FLAG IP RNA to respective total RNA comparisons shown in Fig. 4 and Sup. Fig. 4.

**Supplementary Table 6:** Full DAVID GO term analysis results based on up-regulated differential gene expression values of FLAG IP RNA to respective total RNA comparisons shown in Fig. 4 and Sup. Fig. 4.

**Supplementary Table 7:** Overview of RNAseq experiments, showing analyzed strains and batches in which they were sequenced.

### *C. elegans* strains and culture

All *C. elegans* strains were generated and maintained at 20 °C on nematode growth medium (NGM) agar plates, seeded with the Escherichia coli (*E. coli*) strain OP50-1 (CGC). After CRISPR engineering, all new alleles were outcrossed against the wildtype Bristol N2 strain. Before conducting fitness measures (lifespan, generation time and brood size), collections for biochemistry and RNAseq, all strains were bleached to (*E. coli*) strain OP50. All strains used in this study are listed in Sup. Table 2, including outcrossing information. Genotyping primers used in this study are listed below.

### CRISPR edits to endogenous *rpl-22*

N-terminal 3xFLAG tagged RPL-22 allele *rpl-22(wbm58):*

3x FLAG sequences were amplified out of plasmid pYZ16 using primers *rpl-22* N term flag HR F1

(5’atattttgtaagatctcaatttttactttcaggtaatcatgGACTACAAAGACCATGACG) and *rpl-22* N-term Flag HR R1

(5’gaagagcTttTttggccgacttggcgtgtggctttgggacCTTGTCATCGTCATCCTTGT) followed by a second round of amplification using primers rpl-22 N term flag HR F1 and rpl-22 N term HR R2

(5’tcaacgttgaacttgaggtgcaccttcttcttgcgaagagcTttTttggccgact) to extend the 3’ arm of the homology repair (HR) template sufficiently beyond the *rpl-22* crRNA cut site. A CRISPR mix was made according to Paix *et al*. (2015) and Silva-Garcia *et al*. (2019), and included the HR template, a crRNA targeting the 5’ end of *rpl-22 (rpl-22 5’* crRNA: 5’acacgccaagtcggccaaga), purified Cas9 protein, as well as a co-CRISPR crRNA and HR template to serve as a marker of CRISPR activity (Paix et al., 2015; Silva-Garcia et al., 2019). CRISPR mix was injected to N2 animals on day 1 of adulthood. Progeny from injected animals that showed the most CRISPR activity (as evidenced by a high proportion of animals edited by the co-CRISPR marker) were screened by PCR for presence of the 3x FLAG knock-in. The *rpl-22(wbm58)* allele was identified by PCR, sequenced to ensure a clean edit, and subsequently outcrossed with N2 two times to generate strain MSD470; 6 times with N2 to generate strain WBM1344.

C-terminal tagging of endogenous RPL-22 with 3x FLAG sequences in *rpl-22(wbm53)* [rpl-22::3xflag] and GFP in *rpl-22(wbm85)* [rpl-22::GFP]: CRISPR and CRISPR mixes done as above, using a crRNA specific to the 3’ sequences of endogenous *rpl-22 (rpl-22 3’* crRNA: 5’caaaaaaccgagtttactcg), *dpy-5* crRNA co-CRISPR, and edit-specific HR templates. To generate the HR template for the rpl-22::3xflag edit, 3x flag sequences were amplified from plasmid pYZ16 using primers

rpl-22 3x flag HR F

(5’ttccacatcaacgacggagaggacgcaggaagcgaTcacgagGACTACAAAGACCATGAC)

and

rpl-22 SL2 HR R

(5’aagacaagcagttaactaggtgaaagtaggatgagacagcttactcgtggtcgcttcctg).

To generate the HR template for the rpl-22::GFP edit, GFP cDNA was amplified from lysates of a strain carrying a GFP transgene using the primers rpl-22 GFP HR F (5’ttccacatcaacgacggagaggacgcaggaagcgaTcaTgagAGTAAAGGAGAAGAACTT)

and

GFP rpl-22 3’ UTR HR R

(5’tttgaaacaacaatttattccaacaacaaaaaaccgagtttaTTTGTATAGTTCATCCAT). As above, the *rpl-22(wbm53)* [rpl-22::3xflag] edit was screened for by PCR, sequenced and outcrossed 2 times to generate strain MSD485. The *rpl-22(wbm85)* [rpl-22::GFP] edit was screened for by expression of GFP under a fluorescent dissection scope, confirmed by PCR and sequencing, and outcrossed 3 times to N2 to generate strain WBM1407.

primers used to genotype *rpl-22(wbm58)*:

rpl-22 5’ flag geno F 5’ CGTTTATTCCTGAAGATGCCG

rpl-22 5’ flag geno R 5’ GAGAATTCCATCCTCAACTGG

170 bp N2, 236 bp *rpl-22(wbm58)*

primers used to genotype *rpl-22(wbm53)* and *rpl-22(wbm85)*:

rpl-22 3’ F 5’ cctcaagtaccttaccaaga

rpl-22 3’ UTR R 5’ cattccctactggtactcga

243 bp band in WT, 303 bp band in *rpl-22(wbm53)*, 948 bp band in *rpl-22(wbm85)[rpl-22::GFP]*

### Engineering of SKI TRIP transgenic strains

To generate single copy knock-in alleles driving tissue-specific expression of N-terminal FLAG-tagged RPL-22, full-length *rpl-22* cDNA was amplified from N2 mixed stage cDNA using primers

3x flag rpl-22 HR F

(5’TAAAGATCATGACATCGATTACAAGGATGACGAcGAtAAGgtcccaaagccacacgcc aa) and

rpl-22 SL2 HR R:

(5’aagacaagcagttaactaggtgaaagtaggatgagacagcttactcgtggtcgcttcctg) and knocked into existing and newly engineered SKI LODGE cassettes by CRISPR (Silva-Garcia et al., 2019). The stretches of homology used to flank the *rpl-22* transcript are identical to the 3x flag sequences 5’ of the *dpy-10* edit site in the SKI LODGE cassette and the SL2 trans splicing sequences 3’ of the *dpy-10* edit site, so the same HR template and CRISPR mix was able to be used to edit existing SKI LODGE strains WBM1214 (*N2, wbmIs88[eft-3p::3XFLAG::dpy-10 site::SL2::wrmScarlet::unc-54 3’UTR, *wbmIs67]*) for somatic expression, WBM1215 (*N2, wbmIs89[rab-3p::3XFLAG::dpy-10 site::SL2::wrmScarlet::rab-3 3’UTR,*wbmIs68]*) for neuronal expression, as well as the newly generated strain WBM1325 (*N2, wbmIs114[myo-3p::3XFLAG::dpy-10 site:SL2::wrmScarlet::unc-54 3’UTR, *wbmIs63]*, 6x outcrossed with N2) for expression in muscle. CRISPR mixes were made according to Silva-Garcia et al. (2019) and injected into day 1 adults of the SKI LODGE strains (Silva-Garcia et al., 2019). Candidates were screened from jackpot plates for the presence of the *rpl-22* knock-in by PCR, sequenced, and outcrossed to N2. The resulting SKI TRIP alleles and outcrossed strains are as follows:

Somatic SKI TRIP knock-in: *wbmIs119[eft-3p::3XFLAG::rpl-22::SL2::wrmScarlet::unc-54 3’UTR, *wbmIs88] V* was outcrossed 10 times to N2 to generate strain WBM1364.

Neuronal SKI TRIP knock-in: *wbmIs99[rab-3p::3XFLAG::rpl-22::SL2::wrmScarlet::rab-3 3’UTR, *wbmIs89] IV* was outcrossed 2 times to N2 to generate strain MSD473; 6 times to N2 to generate strain WBM1340.

Muscle SKI TRIP knock-in: *wbmIs118[myo-3p::3XFLAG::rpl-22::SL2::wrmScarlet::unc-54 3’UTR, *wbmIs114] I* was outcrossed 6 times to N2 to generate strain WBM1339.

Intestinal SKI TRIP knock-in: *wbmIs131[nep-17p(mini)::3XFLAG::rpl-22::SL2::wrmScarlet::unc-54 3’UTR, *wbmIs119] V* was outcrossed 6 times to N2 to generate strain WBM1470.

The intestinal SKI TRIP knock-in allele was engineered by excising *eft-3p* sequences from the *eft-3p::3x flag::rpl-22* SKI TRIP cassette in strain WBM1364 using crRNAs targeting sequences 5’ of the *eft-3p* (5’TAAAAGACCAAAGGTGCcgg) and sequences in the 3xFLAG (5’ATGGACTACAAAGACCATGA) and replacing it with 331bp of promoter region upstream of the translational start site of the intestine-specific gene *nep-17* (McGhee et al., 2007). The HR template used to introduce *nep-17p* sequences was amplified from lysates from *C. elegans* strain WBM1450 using primers nep-17p on V

(5’tcaccgtttcgtcgcgtgtcgctcccccgccctaattataagactaatacgtatat) and rpl-22 SL2 HR R (5’aagacaagcagttaactaggtgaaagtaggatgagacagcttactcgtggtcgcttcctg), followed by another round of PCR using primers Chr V SKI HR F universal

(5’ctcacactattcccaaatctagactcatctatcactaatgtcaccgtttcgtcgcgtgtc) and rpl-22 SL2 HR R to extend the HR arms. CRISPR mixes were prepared as above, according to Paix *et al*. (2015) (Paix et al., 2015). Successful edits for the *nep-17p* knock-in were screened for under a fluorescent dissection scope, looking for a switch in wrmScarlet expression from all soma to expression specifically in the intestine. Candidates with intestine-specific expression of wrmScarlet were verified by sequencing, and outcrossed to N2 using fluorescence of wrmScarlet as a marker.

Neuronal SKI TRIP knock-in on Chromosome V: *wbmIs133[rab-3p::3XFLAG::rpl-22::SL2::scarlet::rab-3 3’UTR, V:8645000*wbm127]* was outcrossed 6 times to generate strain WBM1471.

The neuronal SKI TRIP allele on Chromosome V allele was generated by amplifying 3x flag rpl-22 SL2 Scarlet sequences from *C. elegans* lysates from strain WBM1340 using primers 3x flag F universal

(5’ATGGACTACAAAGACCATGACGGTGATTATAAAGATCATGACATCGATTACAAGGATGAC) and

rab-3p 3’ UTR mega R (5’ cgatccatatatctggtgcctagatgttgagagagggaaaactgagagctacgcgc).

The resulting 1540 bp HR template was knocked into a newly generated neuronal SKI LODGE cassette on Chrom V, *wbmIs127[rab-3p::3XFLAG::dpy-10 crRNA::rab-3 3’UTR, V:8645000]* by CRISPR, according to (Silva-Garcia et al., 2019). Progeny from jackpot plates were screened for successful knock-in of *rpl-22::SL2::wrmScarlet* under a fluorescent dissection scope by expression of wrmScarlet in the nervous system. Candidates identified were sequenced for clean knock-in of *rpl-22::SL2::wrmScarlet* into the cassette.

### Microscopy

All strains in this study were imaged on day 1 of adulthood. Adult animals were anesthetized in 0.5 mg/mL tetramisole (Sigma, T1512) diluted in M9 for approximately 15 minutes, and then mounted on 2% agarose pads. Images were acquired on a Zeiss Imager.M2 microscope outfitted with a Colibri 7 Type R[G/Y]CBV-UV Light Source and an AxioCam 506 Mono Camera, and processed using ZEN 3.0 pro software.

### Lifespan assays

Gravid day 1 adults were allowed to lay eggs for ∼4 h. The offspring was used for lifespan analysis. The L4 stage was defined as day 0 and a minimum of 120 *C. elegans* were used per strain and condition. Worms were kept at 20 °C on NGM plates seeded with OP50 *E. coli*. The animals were transferred every day or every second day to fresh plates until they reached the post reproductive stage. Scoring was performed every 1-2 days by monitoring (touch-provoked) movement and pharyngeal pumping. In all lifespan experiments, worms that had undergone internal hatching, vulval bursting, or crawled off the plates were censored. Data were assembled on completion of the experiment. Statistical analyses were performed with the Mantel-Cox log rank method in Prism (Version 8.4.3). All detailed statistics on the lifespan assays and on their repeats are depicted in Sup. Table 1.

### Generation time assays

For generation time assays, synchronized eggs were allowed to develop to adult worms on single NGM plates seeded with OP50 bacteria, until they laid the first egg, which was defined as generation time. After 55-60 h after egg lay, animals were scored every hour for the presence of the first laid egg. 15 animals were assayed per genotype. Error bars represent means ±SD.

### Brood size assays

For brood size assays, the total number of progeny laid by each of the animals assayed for their generation time (above) was counted. The parent animal was transferred to fresh plates every 12-24 h for the first 5 days to facilitate counting. The total number of viable progeny on each transfer plate was counted and summed to get the total number of progeny produced by each of the 15 individual parental animals. Parental animals that burst or died within the first 5 days were censored. Error bars represent means ±SD.

### Polysome profiling

For the analysis of mRNA translation via polysome profiling based on Großhans and Ding (2009), synchronized gravid day 1 adults were grown on NGM plated seeded with OP50 (Ding and Großhans, 2009). Per genotype and replicate, about 12000 worms were harvested and washed twice with M9, once with M9 supplemented with 1 mM cycloheximide (Sigma) and once with lysis buffer (20 mM Tris pH 8.5, 140 mM KCl, 1.5 mM MgCl2, 0.5 % Nonidet P40, 1 mM DTT, 1 mM cycloheximide). Worms were pelleted and resuspended in 350 μL cold lysis buffer, supplemented with 1 % sodiumdeoxycholate (DOC, Sigma). Resuspended worms were lysed using a chilled Dounce homogenizer. Ribonuclease inhibitor RNasin (Promega) was added to samples used for RNA sequencing at a concentration of 0,4 U/μL. Samples were then mixed and incubated on ice for 30 min, followed by a centrifugation step (12000 g, 10 min, 4 °C) for clearance. The pellet was discarded and the RNA concentration of the supernatant was estimated by absorbance measurement at 260 nm. At least 50 µl of lysate were taken aside as corresponding total RNA at this point, diluted in an equal amount of TRI reagent (Zymo Research) and frozen at -80 °C. To prepare sucrose gradients, 15 % (w/v) and 60 % (w/v) sucrose solutions were prepared in basic lysis buffer (20 mM Tris pH 8.5, 140 mM KCl, 1.5 mM MgCl2, 1 mM DTT, 1 mM cycloheximide). Linear sucrose gradients were produced using a Gradient Master (Biocomp). Equivalent amounts of sample (around 400 μg RNA) were loaded onto the gradient and centrifuged at 39000 g for 3 h at 4 °C, using an Optima L-100 XP Ultracentrifuge (Beckman Coulter) and the SW41Ti rotor. To analyze the sample on the gradient during fractionation, absorbance at 254 nm was measured and recorded (Econo UV monitor EM-1, Biorad), using the Gradient Profiler software (version 2.07). Gradient fractionation was performed from the top down using a Piston Gradient Fractionator (Biocomp) and a fraction collector (Model 2110, Biorad). Gradients were fractionated in 20 fractions of equal volume. In an initial experiment, the ribosomal fractions were validated by analyzing RNA from each fraction via agarose gel electrophoresis. The 18 S and 28 S rRNA signals were used as indicators for the 40 S ribosomal subunit, the 60 S ribosomal subunit and fully assembled ribosomes. For more precise analysis of ribosomal fractions, they were collected by hand according to their absorbance profile.

For RNA sequencing experiments, one single fraction comprising all polysomal fractions (corresponding to Figure 1, polysome profiling sequencing A) or one fraction comprising the 60 S ribosomes, the 80 S ribosomes and polysomes (corresponding to Figure 2 and Sup. Figure 2, polysome profiling sequencing B) was collected per sample. RNA extraction from total lysates and polysome fractions was performed using the Direct-zol RNA MicroPrep Kit (Zymo Research) according to the manufacturer’s recommendations.

For Western blot analyses from polysome profiling fractions, protein precipitations from selected fractions were performed using trichloroacetic acid (TCA). Fractions of 600 μL were collected from the polysome profiling. For protein precipitation of each 600 μL fraction, 100 μL of 100 % TCA and 800 μL cold acetone were added. Precipitation was performed at -80 °C overnight. Samples were thawed carefully on ice and centrifuged at 16000 g for 10 min at 4 °C. Precipitates were washed two times with 900 μL cold acetone. Acetone was removed by pipetting and evaporation for 1 h at room temperature. Subsequently, pellets were resuspended in 50 μL of 1x Laemmli Sample Buffer (BioRad) with 2.5 % v/v β-mercaptoethanol, incubated for 10 min with constant shaking on 95 °C and sonicated twice for 2 min before analysis via SDS-PAGE and western blot as described below.

### Western blotting

For Western blotting, respective polysome profiling or IP samples, each with total and elution fractions, were taken to reducing sodium dodecyl sulfatepolyacrylamide gel electrophoresis (SDS-PAGE; Gels: ThermoFisher NuPAGE 4 to 12 %, Bis-Tris). The separated proteins were transferred to nitrocellulose membranes (0.2 µm; Biorad), followed by blocking with milk or bovine serum albumin (BSA) and antibody labelling with specific antibodies to FLAG (Sigma F1804; 1:5000 in 3 % milk) or RPS-6 (Abcam ab70227; 1:1000 in 1 % BSA). Immunolabelling was visualized using chemiluminescence kits (SuperSignal West Pico PLUS Chemiluminescent Substrate; ThermoScientific) on a Chemidoc MP Imaging System (Biorad) and analyzed with the ImageLab Software (version 5.2, Biorad).

### Immuno-precipitation of tagged ribosomes (SKI TRIP experiments)

SKI TRIP experiments were optimized based on previous TRAP models (McLachlan and Flavell, 2019; Sanz et al., 2009). Per SKI TRIP sample, 50 µl Protein G coated dynabeads (ThermoFisher) were washed with 1 ml PBS and incubated with 1 ml PBS and 0.1 μg monoclonal anti-FLAG antibody (Sigma F1804) per μL of dynabeads at 4 °C and constant rotation overnight. Per FLAG IP, 30000 synchronized day 1 worms were collected in M9 buffer, washed two times in M9 and one final time in M9 including 1 mM cycloheximide (Sigma). Samples were then washed with lysis buffer prepared in nuclease free water (10 mM 4-(2-hydroxyethyl)-1-piperazineethanesulfonic acid (HEPES) pH 7.4, 150 mM KCl, 5 mM MgCl2, 0.5 mM DTT, 1 mM cycloheximide, EDTA free Protease Inhibitor). Worms were pelleted at 2000 g for 2 min and the supernatant was discarded. The residual worm pellet (about 500 μL) was shock frozen by dropping it into liquid nitrogen in fractions of approximately 20 μL using a P200 pipette. The resulting frozen worm pearls were carefully lysed using the TissueLyserII (Qiagen) or the CryoMill (Retsch) for twice 1 min at 30 Hz, using metal beads of 5 mm diameter. Keeping all parts and the samples RNase-free and frozen using dry ice and liquid nitrogen at all times was crucial. Lysed worm powder was thawed on ice in a final volume of 3.75 ml lysis buffer including 0.5 % v/v NP-40 (Sigma), 0.4 U/μL RNasin (Promega), 10 mM ribonucleoside vanadyl complex (RVC by NEB), 33 mM 1,2-diheptanoylsn-glycero-3-phosphocholine (DHPC by Merck) and 1 % w/v sodium deoxycholate (Sigma). It is crucial to add the mentioned lysis buffer in a volume to reach approximately 8 worms/μL, taking the starting volume of the ∼500 µl worm pellet into account. In case of varying worm or pellet amounts, the volume of added lysis buffer should be adjusted in this step. Samples were incubated for 30 min on ice. After centrifugation at 12000 g for 10 min at 4 °C, the supernatant was transferred to new tubes (kept on ice at all times) and the pellet was discarded. As corresponding total RNA sample, 100 μL of supernatant was mixed with 100 μL of RLT buffer (Qiagen RNeasy Kit) including 1 % v/v ß-mercaptoethanol. This total RNA sample was then vortexed, incubated for 10 min at room temperature and stored at -80 °C. The remaining supernatant was pre-cleared with washed 10 μL protein G coated Dynabeads (washed once with low-KCl wash buffer: 10 mM HEPES pH 7.4, 150 mM KCl, 5 mM MgCl2, 1 % v/v NP-40) without coupled antibody for at least 1 h. Samples were then incubated with washed FLAG-antibody-coupled beads (washed once with low-KCl wash buffer) for at least 2 h at 4 °C and constant rotation. Beads were washed four times with high-KCl wash buffer (10 mM HEPES pH 7.4, 350 mM KCl, 5 mM MgCl2, 1 % v/v NP-40, 0.04 U/μL RNasin).

For the elution of protein, the beads were incubated with 50 μL of 4x Laemmli Sample Buffer (BioRad) including 2.5 % v/v ß-mercaptoethanol, followed by elution of proteins at 95 °C for 10 min, separation of the eluate from the beads, and analysis by SDS-PAGE and Western blotting.

Alternatively, for RNA elution, the beads were incubated in 350 μL of RLT buffer (Qiagen RNeasy Kit) including 1 % v/v ß-mercaptoethanol for 10 min at room temperature. The eluate was separated from the beads and RNA purification was performed using the Qiagen RNeasy Kit following the manufacturer’s protocol, analyzed using Agilent Technologies 2200 TapeStation System and further processed in RNA sequencing experiments.

### RNA sequencing

For sequencing of RNA obtained from polysome profiling and FLAG IP samples, RNA was eluted and purified as described above in three replicates per experiment. RNA sequencing was performed at the Molecular Biology Core Facilities at the Dana-Farber Cancer Institute (Boston, USA). All RNA samples were processed with a Kapa mRNA library preparation using a starting amount of 200 ng RNA per sample. The RNA libraries were sequenced on an Illumina NoveSeq with the following parameters: at least 20 M reads, 50 bp paired-end sequencing for sample batch AL7663 and 150 bp paired end sequencing for sample batches AL8842 and AL9212. An overview of the sequencing batches can be found in Sup. table 7.

### RNA sequencing analysis

RNA sequencing analysis was performed with the Bioinformatics Core Facility at the Max Planck Institute for Biology of Ageing (Cologne, Germany). Raw reads were mapped to ce11 ENSEMBL build 103 (Howe et al., 2021) (WBcel235, GenBank assembly accession: GCA_000002985.3) using kallisto version 0.46.1 (Bray et al., 2016). Gene counts were quantified in the same step. Differential gene expression between RNA obtained by polysome profiling or FLAG IP and respective total RNA of the same SKI TRIP strain was calculated using DESeq2 version 1.30.1 (Love et al., 2014) in R version 4.0.4.

All principal component analyses (PCAs) were performed on the top 20 % variable genes for each PCA. Genes present in all samples within each respective PCA were analyzed using the R function prcomp. The first and second component were visualized as a scatterplot, indicating the variance depicted by each component on the axis label. For the PCAs in Fig. 1H and Fig. 2D, the principal components were calculated based on DESeq2 normalized ratios per replicate, normalizing polysome profiling RNA or FLAG IP RNA to respective total RNA expression values (FLAG IP RNA/(total RNA + 1)). For the PCAs in Sup. Fig. 2A and Fig. 4A, RNA expression values from total and elution RNA samples from polysome profiling or FLAG IPs per replicate were used. To visualize the samples of different sequencing batches in one PCA (Fig. 4A), ComBat batch correction was used (R package sva version 3.38.0) (Leek et al., 2021).

The scatterplots in Fig. 1I and Fig. 2E show the average ratio of polysome profiling RNA or FLAG IP RNA over respective total RNA expression from three replicates after log2 transformation. These figures show genes identified in both groups. The R function SmoothScatter was used. The Spearman Correlation was calculated using the genes represented in the scatterplot. Darker blue color intensity represents a higher density of genes.

The heatmaps in Fig. 4B and Sup. Fig. 4A were calculated based on DESeq2 normalized ratios per replicate (FLAG IP RNA/(total RNA + 1)), analyzing reference genes known to be expressed in selected tissues. The following genes were selected:

**Germline**: *deps-1* (Zou et al., 2019), *rec-8*(Brenner and Schedl, 2016), *him-8, gld-1* (Brenner and Schedl, 2016), *inx-8* (Gordon et al., 2020)

**Muscle**: *mlc-2* (GuhaThakurta et al., 2004), *unc-89* (Small et al., 2004), *F45D3.2* (Zhao et al., 2007), *jph-1* (Yoshida et al., 2001), *unc-22* (Matsunaga et al., 2017)

**Intestine**: *ges-1* (Zou et al., 2019), *asp-5, K12H4.7, elo-5, T21H3.1* (McGhee et al., 2007)

**Neurons**: *ric-19, sng-1, unc-1* (Ruvinsky et al., 2007), *unc-119, unc-47* (Bessa et al., 2013)

The DAVID GO term analyses in Fig. 4C-F and Sup. Fig. 4B were calculated based on up- or down-regulated differential gene expression values of FLAG IP RNA to respective total RNA comparisons from three replicates per SKI TRIP strain. The analyses were performed using the AGEpy library version 0.8.2 (Metge et al., 2018) (Huang da et al., 2009) in Python version 3.7. They were visualized using FLASKI (Iqbal et al., 2021).

## Acknowledgements

We thank all Denzel and Mair laboratory members for helpful discussions. We deeply thank F. Metge, J. Boucas and all members of the Bioinformatics Core Facility at the MPI for Biology of Ageing (Cologne, Germany) for their great support in all bioinformatic analyses throughout this manuscript. We thank the Molecular Biology Core Facilities at the Dana-Farber Cancer Institute (Boston, USA) for sequencing. Figures 1A, 1F and 2A were created with BioRender.com.

## Author contributions

M.S.D., W.B.M., L.E.W., A.L. and E.H.W.B. designed the study. A.L. designed and generated the SKI TRIP alleles used in this study. C.H. designed and generated *wbmIs127*. L.E.W. and E.H.W.B. performed biochemical experiments and generated samples for RNA sequencing. A.L., L.E.W and M.C.P-M. performed fitness measures on all *C. elegans* strains. A.L., L.E.W., and E.H.W.B. wrote the manuscript. L.E.W., A.L., M.S.D. and W.B.M. finalized the study and the manuscript.

---

**Sup. Figure 1:**
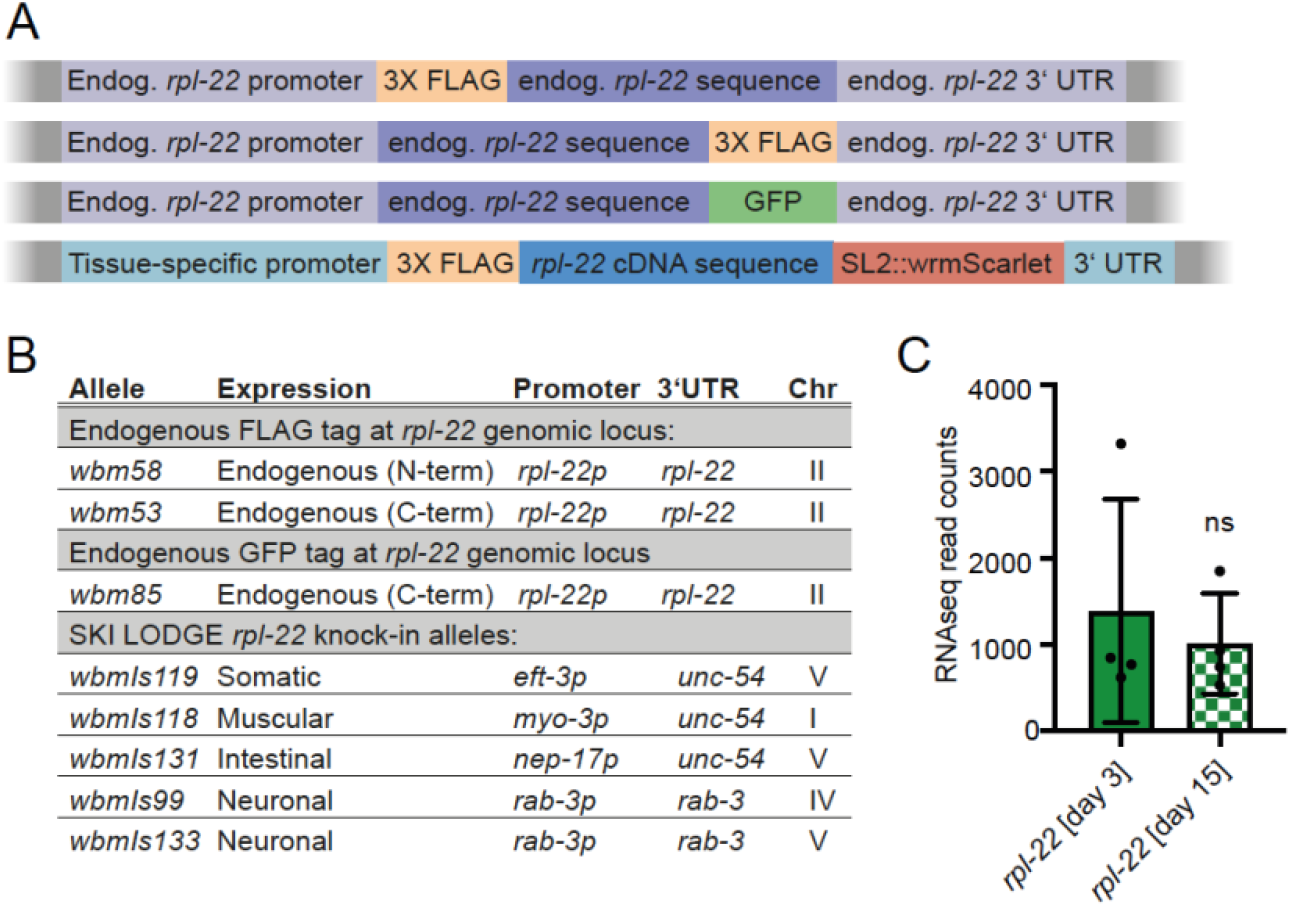
Design of endogenous and tissue-specific FLAG::RPL-22 SKI TRIP strains. (**A**) Schematic overview of the genetic SKI TRIP constructs. Upper 3 panels: CRISPR/Cas9 engineered FLAG- or GFP-tags at the endogenous *rpl-22* locus. Lower panel: CRISPR/Cas9 engineered single-copy knock-in cassette including an N-terminal FLAG-tag, the *rpl-22* cDNA sequence and the wrmScarlet sequence separated by SL2 trans-splicing signal sequences. The cassette is under defined tissue-specific promotors and selected 3’UTRs (see **B**). (**B**) Detailed information for all SKI TRIP strains. (**C**) mRNA levels of *rpl-22* in wildtype animals at day 3 and day 15 of adulthood, measured by RNAseq (Heintz et al., 2017), depicted as read counts from n=4 experiments (error bars represent means ±SD; unpaired two-tailed t-test; no significant changes were detected). See Sup. Table 2 for full worm strain information.

**Sup. Figure 2:**
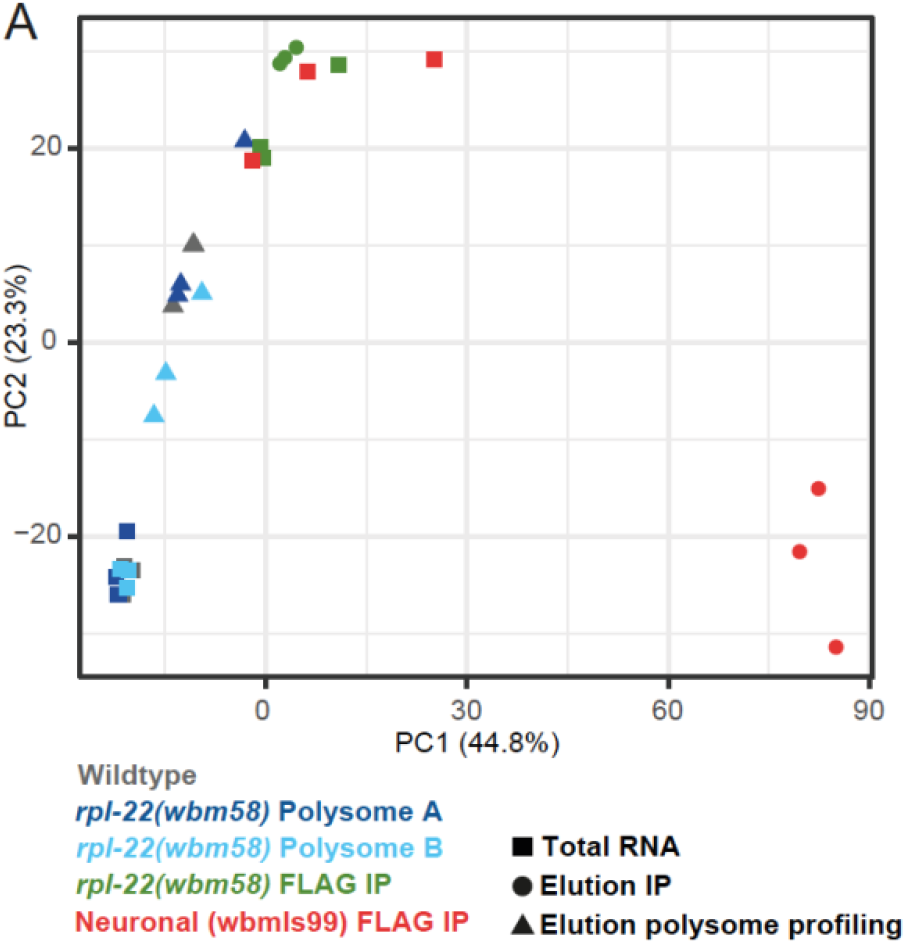
Polysome profiling and FLAG co-IP experiments from endogenous SKI TRIP knock-in strain capture different pools of mRNAs in total and elution fractions. (**A**) PCA using RNA expression values, comparing total and elution RNA from Polysome Profiling or FLAG co-IPs of *rpl-22(wbm58) C. elegans* samples. As reference, also data corresponding to total and elution RNA from FLAG co-IPs of neuronal (*wbmIs99*) SKI TRIP worm samples are included. Data corresponding to n=3 independent replicates from each strain/experiment are shown. Dots represent total RNA, triangles represent elution RNA. PC = Principal Component. See Sup. Figure 1 for detailed information on SKI TRIP strains. See Sup. Table 3 for differential gene expression tables corresponding to the RNA sequencing experiments.

**Sup. Figure 3:**
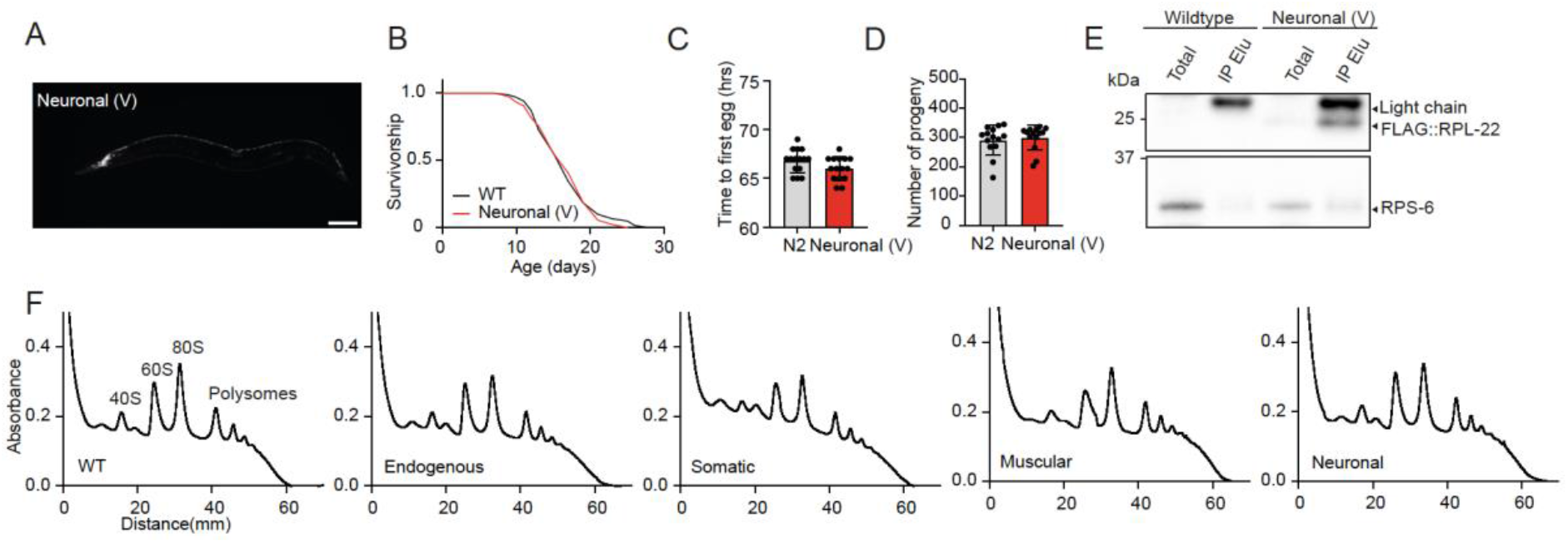
Neuronal SKI TRIP cassette on Chromosome V does not impact overall C. elegans physiology and is sufficient for FLAG co-IPs. Analysis of neuronal (Chr V) SKI TRIP animals (*wbmIs133*). FLAG-tagged RPL-22 is expressed under the neuron-specific *rab-3* promotor. The expression cassette is carrying a Scarlet fluorophore sequence after the FLAG::RPL-22; an SL2 sequence induces trans-splicing of the fluorophore from FLAG::RPL-22. As comparison, the neuronal SKI TRIP strain *wbmIs99* shown in main Figure 3 carries the expression cassette on Chromosome IV. (**A**) Fluorescence microscopy image of a day 1 neuronal (V) SKI TRIP animal. Scale bar = 100 µm. (**B**) Survival of neuronal (V) SKI TRIP animals compared to WT (representative data from n=3 independent experiments). (**C**) Generation time of WT and neuronal (V) SKI TRIP animals (error bars represent means ±SD; representative data from n=2 independent experiments with 15 animals per genotype). (**D**) Brood size of WT and neuronal (V) SKI TRIP animals as total number of viable progeny per individual parental animal (error bars represent means ±SD; representative data from n=2 independent experiments with ≥14 animals per genotype). (**E**) Western blot of day 1 WT and neuronal (V) SKI TRIP samples after FLAG co-IP. Western blot using anti-FLAG and anti RPS-6 antibodies. co-IP total and elution (IP elu) fractions are shown. (**F**) Polysome profiling graphs of day 1 WT and endogenous and selected tissue-specific SKI TRIP samples. “Neuronal (V)” = neuron-specific SKI TRIP *wbmIs133* [WBM1471] animals. “Somatic” = soma-specific SKI TRIP *wbmIs119* [WBM1364] animals. “Muscular” = muscle-specific SKI TRIP *wbmIs118* [WBM1339] animals. “Neuronal” = neuron-specific SKI TRIP *wbmIs99* [MSD473] animals. See Sup. Figure 1 for detailed information on SKI TRIP strains. See Sup. Table 1 for lifespan statistics.

**Sup. Figure 4:**
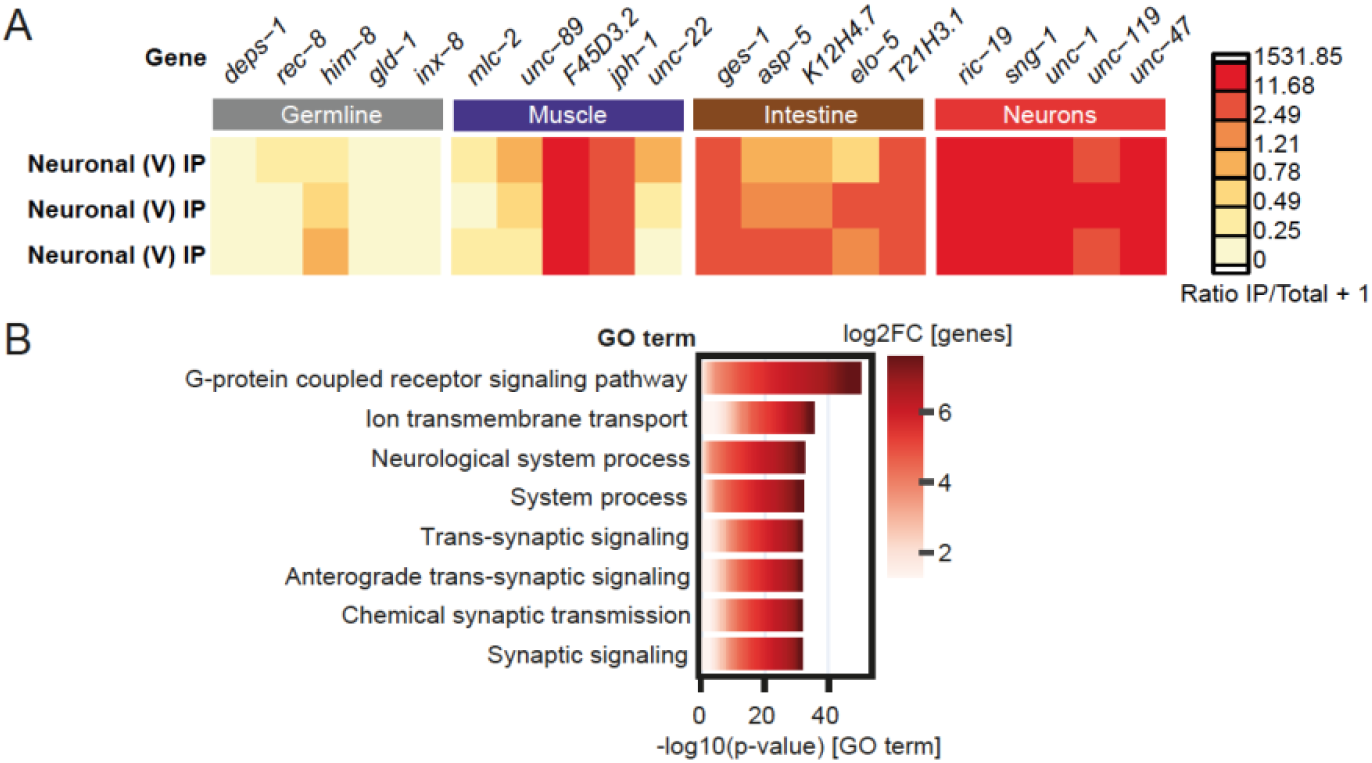
FLAG co-IP efficiently pulls down neuronal mRNAs from samples generated using the neuronal SKI TRIP cassette on chromosome V. FLAG IPs were performed on samples from neuron-specific (Chr V) SKI TRIP allele *wbmIs133*, n=3 independent replicates. Total and eluted RNA were sequenced. (**A**) Heat map is showing expression values of FLAG co-IP elution samples that were normalized to respective total RNA samples from the neuronal (V) SKI TRIP strain. Data shown for selected genes that are known to be expressed in specific tissues as indicated in the figure. According to Wormbase (WS282), the genes *F45D3.2* and *jph-1* are reported to be expressed in muscle and neurons. (**B**) DAVID GO term analysis of mRNAs significantly changed in translation (IP elution RNA values normalized to respective total RNA values of the same strain, using data of n=3 independent replicates). Analyzed were significantly up-regulated transcripts from the neuronal (V) SKI TRIP strain. Shown are the top 8 significant GO terms. “Neuronal (V)” = neuron-specific SKI TRIP *wbmIs133* [WBM1471] animals. See Sup. Figure 1 for detailed information on SKI TRIP worm strains. See Sup. Table 4 for differential gene expression tables corresponding to the RNA sequencing experiments. See Sup. Table 5 and 6 for full DAVID GO term analysis results.

## Notes

### Competing Interest Statement

The authors have declared no competing interest.

